# Behavioral flexibility in an OCD mouse model: Impaired Pavlovian reversal learning in SAPAP3 mutants

**DOI:** 10.1101/435172

**Authors:** Bastijn J. G. van den Boom, Adriana H. Mooij, Ieva Misevičiūtė, Damiaan Denys, Ingo Willuhn

## Abstract

Obsessive-compulsive disorder (OCD) is characterized by obsessive thinking, compulsive behavior, and anxiety, and is often accompanied by cognitive deficits. The neuropathology of OCD involves dysregulation of cortical-striatal circuits. Similar to OCD patients, SAPAP3 knockout mice 3 (SAPAP3^−/−^) exhibit compulsive behavior (grooming), anxiety, and dysregulated cortical-striatal function. However, it is unknown whether SAPAP3^−/−^ display cognitive deficits and how these different behavioral traits relate to one another. SAPAP3^−/−^ and wild-type littermates (WT) were trained in a Pavlovian conditioning task pairing the delivery of visual cues with that of sucrose solution. After mice learned to discriminate between a reward-predicting conditioned stimulus (CS+) and a non-reward stimulus (CS−), contingencies were reversed (CS+ became CS− and vice versa). Additionally, we assessed grooming, anxiety, and general activity. SAPAP3^−/−^ acquired Pavlovian approach behavior similarly to WT, albeit less vigorously and with a different strategy. However, unlike WT, SAPAP3^−/−^ were unable to adapt their behavior after contingency reversal, exemplified by a lack of re-establishing CS+ approach behavior (sign tracking). Surprisingly, such behavioral inflexibility, decreased vigor, compulsive grooming, and anxiety were unrelated. This study demonstrates that SAPAP3^−/−^ are capable of Pavlovian learning, but lack flexibility to adapt associated conditioned approach behavior. Thus, SAPAP3^−/−^ do not only display compulsive-like behavior and anxiety, but also cognitive deficits, confirming and extending the validity of SAPAP3^−/−^ as a suitable model for OCD. The observation that compulsive-like behavior, anxiety, and behavioral inflexibility were unrelated suggests a non-causal relationship between these traits and may be of clinical relevance for OCD patients.

## INTRODUCTION

Obsessive-compulsive disorder (OCD) is a psychiatric disorder that is characterised by recurrent unwanted thoughts, anxiety, and compulsive behavior, but is also often associated with cognitive deficits (1–5). The persistence of maladaptive patterns of inflexible thoughts and behavior suggest a lack of cognitive flexibility (4), the ability to adapt behavior in response to changing situational requirements. Consistently, OCD patients were found to show abnormalities in cognitive flexibility (6–10).

Preclinical animal models are a valuable tool to elucidate neurobiological mechanisms of OCD, but also promise to unravel how different symptoms relate to one another. Evidence implicates dysregulation of projections from cortex to striatum in the neuropathology of OCD (11–14). Mice with genetic deletion of Synapse-associated protein 90/postsynaptic density protein 95 associated protein 3 (SAPAP3^−/−^), a postsynaptic scaffolding protein predominantly expressed in cortico-striatal circuits (15–17), exhibit dysregulation of the same pathway (16, 17). SAPAP3^−/−^ display compulsive-like grooming that can be decreased by deep-brain stimulation of cortico-striatal pathway associated areas (18) and optogenetic stimulation of cortico-striatal projections (19). Conversely, stimulation of cortico-striatal projections in wild-type mice evokes lasting increases in grooming (20). In addition to excessive grooming, SAPAP3^−/−^ mice show increased anxiety, both of which can be reduced by viral rescue of striatal SAPAP3 (16). Similarly, administration of selective serotonin reuptake inhibitors, the primary pharmacotherapy for OCD, normalize self-grooming and anxiety-like behavior in SAPAP3^−/−^ (16).

Despite this promising validation of the model, cognitive deficits have not been assessed in SAPAP3^−/−^ until now (this manuscript and (21)). To study cognitive flexibility in both humans and animals, reversal learning paradigms are often used (22, 23). Abnormal reversal learning performance or altered neural activity during reversal learning is prevalent in OCD patients (6–10). During reversal learning, previously acquired contingencies of stimulus-reward associations are reversed, and the subjects’ adaptation to this is assessed.

Pavlovian conditioning is the most basic type of associative learning, during which a conditioned stimulus (CS) can trigger approach behavior, a procedure called “autoshaping” (24, 25). Autoshaping enables differentiation between approach towards the predictive CS itself (so-called sign tracking), thought to be driven by model-free strategies, and the reward location (goal tracking), presumably driven by model-based strategies (26–28), thereby probing cognitive mechanisms underlying the behavior.

To investigate cognitive flexibility in SAPAP3^−/−^, we trained mice in an autoshaping paradigm in touchscreen boxes. Upon task acquisition, reward contingencies were reversed. In addition, we investigated the relationship between behavioral flexibility, compulsive-like behavior, and anxiety. Such a multi-faceted behavioral investigation of SAPAP3^−/−^ may contribute to the understanding of behavioral deficits in OCD patients.

## MATERIALS AND METHODS

### Subjects

Animal procedures were in accordance with European and Dutch laws and approved by the Animal Experimentation Committee of the Royal Netherlands Academy of Arts and Sciences. Male and female SAPAP3^−/−^ (mean age 9 months, n=20, two excluded due to skin lesions) and wild-type littermates (WT) (mean age 8 months, n=14) were housed individually on a reversed light-dark cycle (lights on from 19:00 to 07:00) and food-restricted to 85% of their free-feeding body weight. All behavioral procedures were performed during the dark phase.

### Procedure

First, grooming was assessed in an open field (OF; pre-autoshaping) for 60 minutes. Next, animals were tested in the autoshaping task, followed by a second 60-minute OF test (post-autoshaping), and 10 minutes on the elevated plus maze (EPM) to probe anxiety. A Janelia Automatic Animal Behavior Annotator classifier (29) was used to quantify self-grooming (see (30)). More methodological details can be found in the supplemental information.

### Pavlovian conditioning (autoshaping)

#### Apparatus

Training was performed in trapezoid-shaped Bussey-Saksida touchscreen chambers (Campden Instruments, Leics, UK) (31) (Fig. 1a). CS appeared in two different positions on the touchscreen (white rectangles presented for 10 s), but were otherwise indistinguishable. Chambers were equipped with infrared beam detectors – one for trial initiation (opposite to touchscreen), two measuring stimuli approaches, and one to count reward magazine entries.

**Figure 1.**
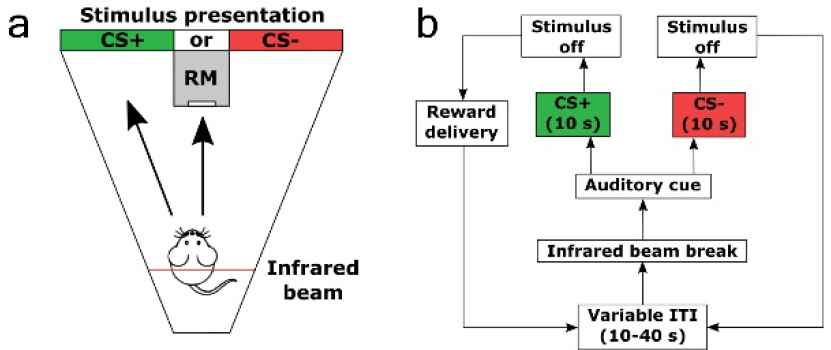
Pavlovian conditioning (autoshaping) in touchscreen chambers. Panel (a) depicts a schematic of top-down view of touchscreen chamber. Illumination of different parts of the screen served as conditioned stimuli (CS). CS+ on the left (green) or CS− on the right (red) (counterbalanced). Besides position, CS were indistinguishable. Approaches (arrows) to CS or reward magazine (RM) were recorded. Panel (b) depicts a schematic of the autoshaping task. Mice initiated trials by interrupting the infrared beam. Left side shows a CS+ trial and right side a CS− trial. Sucrose solution was delivered into the RM 10 s after CS+ onset. Contingency reversal consisted of switching CS (CS+ became CS− and vice versa). CS=conditioned stimulus; RM=reward magazine; ITI=inter-trial interval.

#### Training

Mice were trained twice a day in the autoshaping task for 30 minutes per session, for a total of 72 sessions (36 before and 36 after reversal). A trial started when the mouse interrupted the infrared beam at the back of the chamber after a variable inter-trial interval of 25 s, followed by an auditory cue and visual stimulus presentation on either the left (rewarded conditioned stimulus; CS+) or right (non-rewarded conditioned stimulus; CS−) side of the screen (position counterbalanced between animals). Upon CS+ offset, reward was delivered to the magazine (Fig. 1b).

#### Reversal training

Reward contingencies were reversed after 36 sessions (spatial reversal of CS), whereby the previous CS+ became the CS− and the previous CS− became the CS+.

#### Exclusion criteria

We excluded animals that did not associate the CS+ with reward (32–34). Thus, animals that failed to approach screen or reward magazine during CS+ in over 70% of trials (in the last 10 sessions before reversal) and/or failed to avoid screen or reward magazine during CS− less than 70% of trials in the same sessions, were excluded from the analysis. This resulted in three excluded WT and six excluded SAPAP3^−/−^. Additionally, on a session-by-session basis, individual sessions in which animals initiated less than 10 trials were excluded.

#### Performance measures

We measured the following variables: 1) number of trials initiated, 2) number of trials with an approach towards screen, 3) number of trials with an approach towards reward magazine, 4) combined number of trials with an approach toward screen and/or reward magazine [CS screen approach + CS reward magazine approach], 5) difference (score) of combined number of trials [combined CS+ approaches – combined CS− approaches], and 6) general activity during autoshaping measured as total number of infrared beam breaks outside of CS presentation. Combined number of trials with an approach (during CS presentation) consisted of the animals’ approach to the cue, the reward magazine, or both (24).

### Data analysis

Parametric analyses were used if data were normally distributed, otherwise non-parametric alternatives. *P* values were adjusted for multiple comparisons using the Holm-Bonferroni correction (35). Statistical significance was defined as *p*<0.05.

## RESULTS

### SAPAP3^−/−^ mice phenotyping

Grooming was assessed in the OF before (pre-autoshaping) and after (post-autoshaping) Pavlovian conditioning. A main effect of genotype on grooming duration (F_(1,29)_=8.69, *p*=*.006*), but no main effect of session (F_(1,29)_=0.05, *NS*), nor an interaction effect (F_(1,29)_=0.26, *NS*) was found. SAPAP3^−/−^ showed increased grooming duration both pre-autoshaping (Fig. 2a, mean±SEM: 159.43±19.67 s WT vs. 320.53±58.67 s SAPAP3^−/−^, t_(19.50)_=−2.60, *p*=*0.017*) and post-autoshaping (Fig. 2a, 114±15.71 s WT vs. 312.29±66.62 s SAPAP3^−/−^, U=57, z=−2.46, *p*=*0.026*). Grooming bouts showed a similar effect (supplementary Fig. 1a).

**Figure 2.**
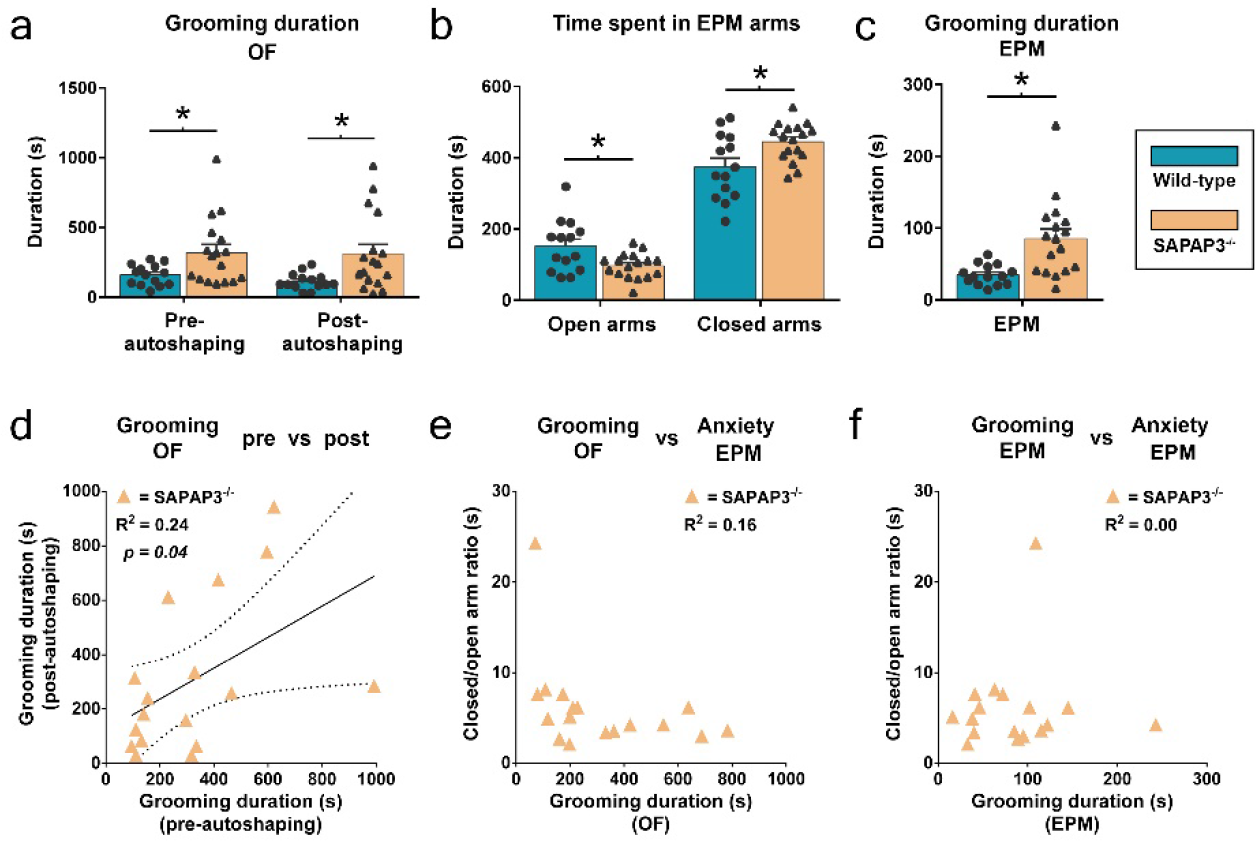
Increased grooming (open field (OF)) and anxiety (elevated plus maze (EPM)) in SAPAP3^−/−^ are not correlated. Grooming duration in the open field before and after Pavlovian conditioning (autoshaping) for wild-type littermates (n=14, blue, circles are individual animals) and SAPAP3^−/−^ (n=17, orange, triangles are individual animals) (a). Time spent in open and closed arms on the EPM as a measure of anxiety (b). Grooming duration during EPM testing (c). Significant correlation between grooming duration in the open field before and after Pavlovian conditioning (d). No correlation between (averaged) grooming duration in the open field and anxiety on the EPM (ratio of time spent in closed/open arms) (e). No correlation between grooming on the EPM and anxiety on the EPM (f). Data are mean+SEM; * p<0.05. OF=open field, EPM=elevated plus maze.

A significant correlation was found between pre-autoshaping and post-autoshaping grooming for SAPAP3^−/−^ (Pearson *R^2^* of 0.243; *p*=*0.04*; 95% CI 0.02-0.79), indicating relatively consistent grooming over time (three months) (Fig. 2d). These grooming durations were averaged to compute a *grooming trait value* per animal.

On the EPM, SAPAP3^−/−^ spent less time in open arms (Fig. 2b: 152.25±19.52 s WT vs. 97.38±8.51 s SAPAP3^−/−^, t_(17.88)_=2.58, *p*=*0.019*) and more time in closed arms compared to WT (Fig. 2b: 374.58±24.27 s WT vs. 445.28±12.96 s SAPAP3^−/−^, t_(20.14)_=−2.57, *p*=*0.036*). Transitions to open and closed arms displayed a similar effect (supplementary Fig. 1b). Similar to the OF test, SAPAP3^−/−^ showed increased grooming duration (Fig. 2c: 35±3.78 s WT vs. 85.53±13.23 s SAPAP3^−/−^, t_(29)_=−3.37, *p*=*0.002*) during EPM testing. Grooming bouts during EPM tested showed a similar effect (supplementary Fig. 1c).

To study the relation between grooming and anxiety, the grooming trait value was correlated with EPM performance (ratio of time spent in closed versus open arms), resulting in no significant correlation (Pearson *R^2^* of 0.16; *NS*; 95% CI −0.74-0.11) (Fig. 2e). In addition, no correlation was found between grooming during EPM testing and EPM performance (ratio of time spent in closed versus open arms) (Pearson *R^2^* of 0.004; *NS*; 95% CI −0.43-0.53) (Fig. 2f).

### General activity during behavioral tasks

Throughout different behavioral tasks, we assessed general activity. Visual inspection of trajectories of SAPAP3^−/−^ in the OF revealed similar movement patterns compared to WT (Fig. 3a, b). During OF testing, movement was quantified during periods when animals did not groom (to exclude effects of grooming on activity). A main effect of genotype on movement was found (F_(1,29)_=14.47, *p*=0.001), but no main effect of time between OF sessions (F_(1,29)_=0.62, *NS*), nor an interaction effect (F_(1,29)_=0.12, *NS*). Post-hoc analyses revealed that SAPAP3^−/−^ showed decreased movement during pre-autoshaping OF (Fig. 3c: 21 081.5±4,001 cm WT vs. 10 644.6±769.1 cm SAPAP3^−/−^, U=38, z=−3.22, *p*=*0.001*) and post-autoshaping OF (Fig. 3c: 21 772.6±2,083.3 cm WT vs. 12 445.2±961 cm SAPAP3^−/−^, U=18, z=−4.01, *p*<*0.0001*). To test if decreased general activity in SAPAP3^−/−^ was due to excessive grooming, movement was divided by time not spent grooming. Similar to movement, a main effect of genotype on movement per minute was found (F_(1,29)_=12.32, *p*<*0.0001*), but no main effect of time in between OF testing (F_(1,29)_=0.51, *NS*), nor an interaction effect (F_(1,29)_=0.16, *NS*). Post-hoc tests revealed that SAPAP3^−/−^ demonstrated decreased movement per minute pre-autoshaping (Fig. 3d: 368.45±71.42 cm/min WT vs. 194.29±13.1 cm/min SAPAP3^−/−^, U=46, z=−2.9, *p*=*0.003*) and post-autoshaping (Fig. 3d: 377.05±37.63 cm/min WT vs. 224.51±15.13 cm/min SAPAP3^−/−^, U=19, z=−3.97, *p*<*0.0001*). Movement during OF was not related to grooming (Pearson *R^2^* of 0.03; *NS*; 95% CI −0.60-0.34) (Fig. 3e).

**Figure 3.**
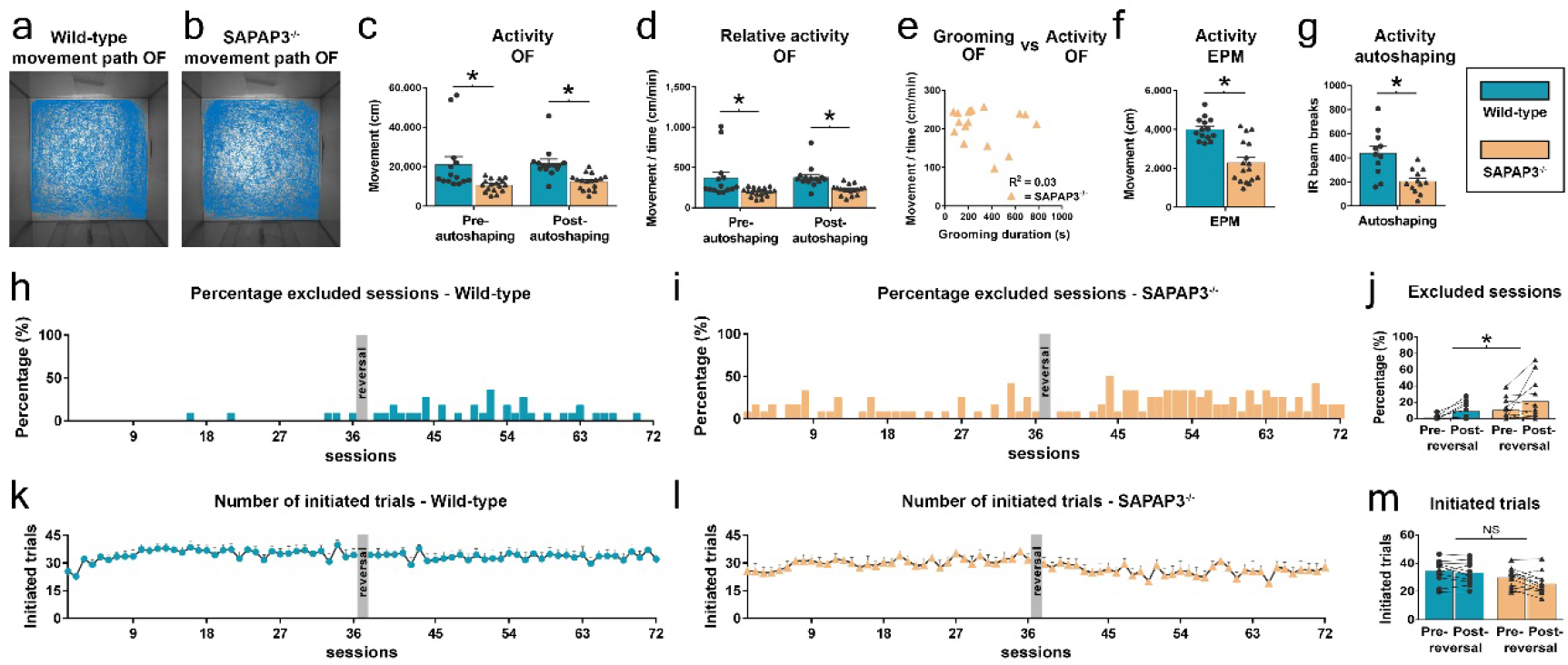
SAPAP3^−/−^ display decreased general activity during behavioral testing that did not affect the overall number of initiated trials. Representative open field (OF) trajectories of a wild-type littermate (WT) (a) and a SAPAP3^−/−^ (b) during OF testing. Activity in the OF measured during periods when animals were not grooming for WT (n=14, blue, circles are individual animals) and SAPAP3^−/−^ (n=17, orange, triangles are individual animals) (c). Average activity in the OF per minute of time *not* spent grooming (d). No correlation between averaged grooming duration and average activity in the OF per minute (e). General activity on the elevated plus maze (f). Beam breaks measured during autoshaping inter-trial intervals (no CS) as a proxy for activity for WT (n=11, circles) and SAPAP3^−/−^ (n=12, triangles) (g). Percentage of WT autoshaping sessions excluded based on criterion of a minimum of 10 initiated trials per session (h). Percentage of SAPAP3^−/−^ autoshaping sessions excluded (i). The average number of excluded WT and SAPAP3^−/−^ sessions is significantly different (j). Average number of initiated trials per session for WT (excluded sessions removed) (k). Average number of initiated trials per session for SAPAP3^−/−^ (excluded sessions removed) (l). Average number of trials initiated by WT and SAPAP3^−/−^ before and after reversal is not significantly different (m). Data are mean+SEM; * p<0.05. OF=open field; EPM=elevated plus maze; NS=not significant.

During EPM testing, SAPAP3^−/−^ exhibited decreased movement (Fig. 3f: 4 008.17±156.34 cm WT vs. 2 292.15±259.57 cm SAPAP3^−/−^, t_(29)_=5.36, *p*<*0.0001*) that was not due to increased grooming (grooming duration EPM versus movement EPM; Pearson *R^2^* of 0.03; *NS*; 95% CI −0.61-0.34).

During autoshaping, average total beam breaks during inter-trial intervals (as a proxy for activity) were averaged across sessions. SAPAP3^−/−^ showed decreased activity compared to WT (Fig. 3g: 438.35±57.82 breaks WT vs. 206.44±28.13 breaks SAPAP3^−/−^, t_(21)_=3.71, *p*=*0.001*).

Mice had to initiate trials by interrupting an infrared beam located opposite to the screen. Some animals initiated only approximately five trials per session, whereas the majority of the animals initiated about 30. Sessions with less than 10 initiated trials were excluded from analyses (Fig. 3h, i, j). Analyses on the number of excluded sessions revealed a significant main effect of genotype (F_(1,23)_=4.26, *p*=*.05*) and a main effect of reversal (F_(1,23)_=7.94, *p*=*.01*), but no interaction (F_(1,23)_=0.19, *NS*), suggesting that although reversal had an effect on trial initiation in both genotypes, SAPAP3^−/−^ were more inactive in general.

After exclusion of inactive sessions, the number of initiated trials for WT revealed no significant effect of reversal (F_(1,569.59)_=2.14, *NS*), nor an effect of session (F_(35,30.3)_=0.73, *NS*) nor an interaction effect (F_(35,30.3)_=1.01, *NS*) (Fig. 3k). SAPAP3^−/−^ displayed a minor decrease in initiated trials after reversal (F_(1,481.69)_=28.31, *p*<*.001*), but no effect of session (F_(35,27.45)_=0.45, *NS*) nor an interaction effect (F_(35,27.45)_=1.01, *NS*) (Fig. 3l). Direct comparison of initiated trials between WT and SAPAP3^−/−^ revealed no difference (Fig. 3m: 33.82±2.39 trials WT vs. 27.83±2.01 trials SAPAP3^−/−^, t_(21)_=1.93, *NS*).

### Autoshaping performance

During CS+ presentation before reversal, WT interacted with the reward magazine (Fig. 4a) as well as the CS itself (Fig. 4c) with no systematic preference. After reversal, WT re-acquired the new reward contingencies, demonstrated by increased CS approaches, but refrained from magazine approaches.

**Figure 4.**
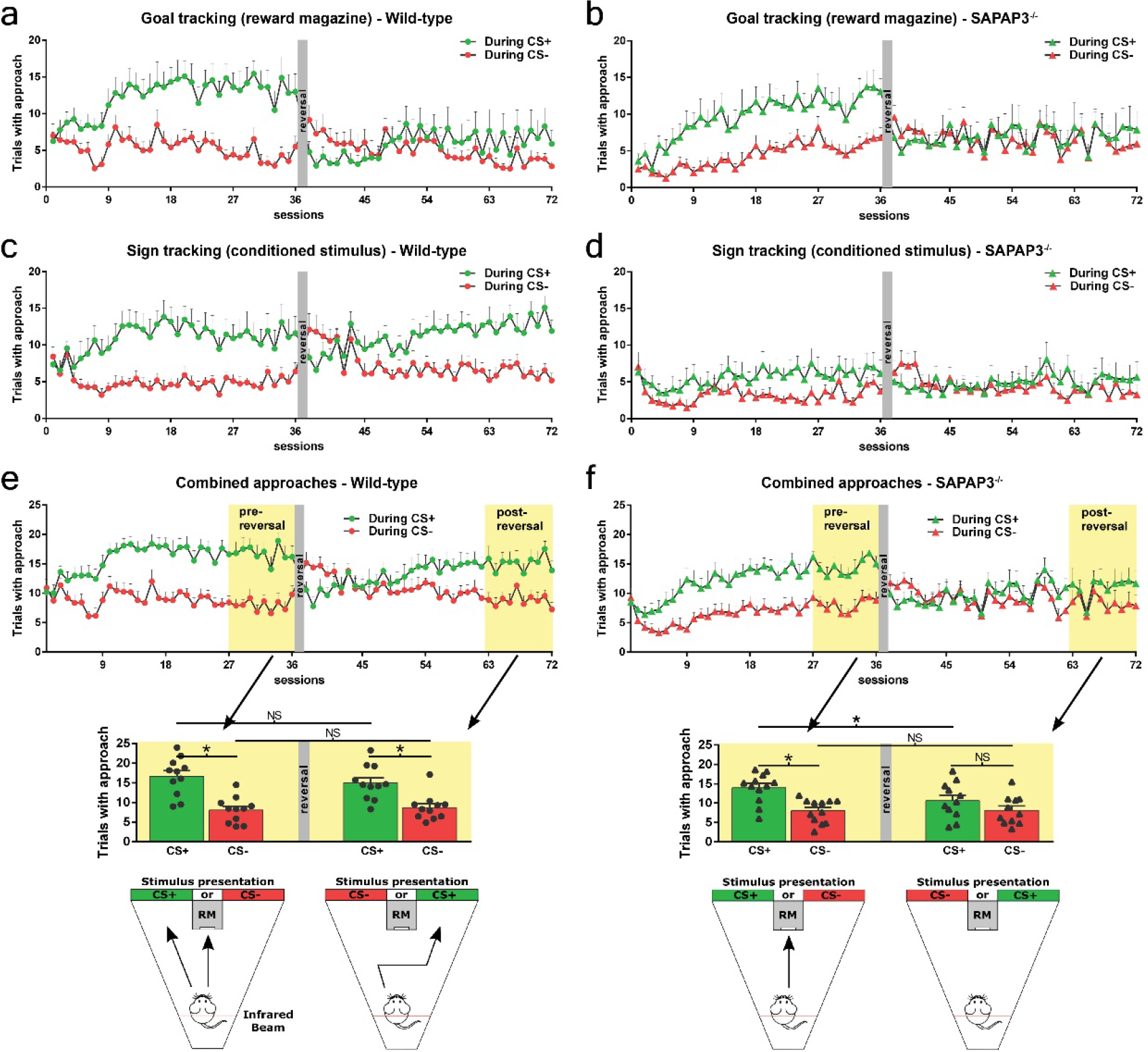
SAPAP3^−/−^ show reduced approaches towards the CS+ (sign tracking) and impaired overall Pavlovian responding after contingency reversal. Number of trials with reward magazine approach (goal tracking) during CS+ (green) and CS− (red) for wild-type littermates (WT) (n=11, circles) (a). Number of trials with reward magazine approach for SAPAP3^−/−^ (n=12, triangles) (b). Number of trials with CS approach (sign tracking) for WT (c). Number of trials with CS approach for SAPAP3^−/−^ (d). Panel (e) depicts the number of trials with at least one approach [CS approach and/or reward magazine approach] for WT. Yellow background indicates the last 10 sessions before reversal and after reversal used for statistical analyses. Bar graphs below show average performance with individual WT depicted as circles. Schematics on bottom display preferred strategy employed by WT before and after reversal. Panel (f) depicts the combined approaches for SAPAP3^−/−^. Bar graphs below show average performance with individual SAPAP3^−/−^ depicted as triangles. Schematics on bottom show preferred strategy before and after reversal. Data are mean+SEM; * p<0.05. CS=conditioned stimulus; RM=reward magazine; NS=not significant.

Similar to WT, SAPAP3^−/−^ learned to discriminate between CS+ and CS−, but mainly only approached the reward magazine (Fig. 4b) and not towards the CS (Fig. 4d). After reversal, SAPAP3^−/−^ showed diminished discrimination between the CS+ and CS−, but still retrieved rewards.

Because mice interacted with both screen and magazine, we calculated combined approaches towards screen and magazine during either CS+ or CS− presentation. This allowed direct comparison of autoshaping performance within and between genotypes, independent of applied behavioral strategy (Fig. 4e, f).

Statistics were performed on the 10 sessions before reversal and on the last 10 sessions after reversal averaged over animals, as performance became asymptotic. In WT, a main effect of CS on autoshaping performance (F_(1,10)_=142.66, *p*<*.0001*), no main effect of reversal (F_(1,10)_=0.21, *NS*), nor an interaction effect (F_(1,10)_=0.11, *NS*) was found. Post-hoc analyses revealed a significant difference between approach behavior towards CS+ and CS− before (Fig. 4e, bar graphs pre-reversal: 16.64±1.46 trials with approaches CS+ vs. 8.07±0.99 trials with approaches CS−, z=−2.93, *p*=*0.009*) and after reversal (post-reversal: 15.02±1.31 trials with approaches CS+ vs. 8.71±0.99 trials with approaches CS−, t_(10)_=9.1, *p*<*0.001*), accompanied by no difference between CS+ before and after reversal (t_(10)_=1.29, *NS*) nor between CS− before and after reversal (t_(10)_=−0.51, *NS*), suggesting that mice reached similar performance after reversal.

In SAPAP3^−/−^, we found a significant main effect of CS on autoshaping performance (F_(1,10)_=33.96, *p*<*.0001*), no main effect of reversal (F_(1,10)_=1.66, *NS*), but accompanied by a significant interaction effect (F_(1,10)_=9.81, *p*=*.01*), suggesting that SAPAP3^−/−^ approached CS+ and CS− differently before and after reversal. Indeed, post-hoc analyses revealed that SAPAP3^−/−^ differentiated between CS+ and CS− before reversal (Fig. 4f bar graphs: 13.99±1.1 trials with approaches CS+ vs. 8±0.88 trials with approaches CS−, t_(11)_=7.96, *p*<*0.0001*), but not after reversal (10.69±1.4 trials with approaches CS+ vs. 8.11±1.11 trials with approaches CS−, t_(10)_=2.53, *p*=*NS*). A significant decrease in CS+ approach behavior was found after reversal (t_(10)_=2.91, *p*=*.045*), but not in CS− approach behavior (t_(10)_=−0.21, *NS*).

### Direct performance comparison between genotypes

We computed a difference score of combined approach behavior for both genotypes (Fig. 5a) and performed statistics on the first 10 sessions before and after reversal (pre-acquisition and post-acquisition, respectively) and the last 10 sessions before and after reversal (pre-maintenance and post-maintenance, respectively). A main effect of reversal on acquisition (F_(1,21)_=96.37, *p*<*.0001*) was found, but no main effect of genotype (F_(1,21)_=0.04, *NS*), nor an interaction effect (F_(1,21)_=0.001, *NS*), suggesting similar acquisition rate between WT and SAPAP3^−/−^.

**Figure 5.**
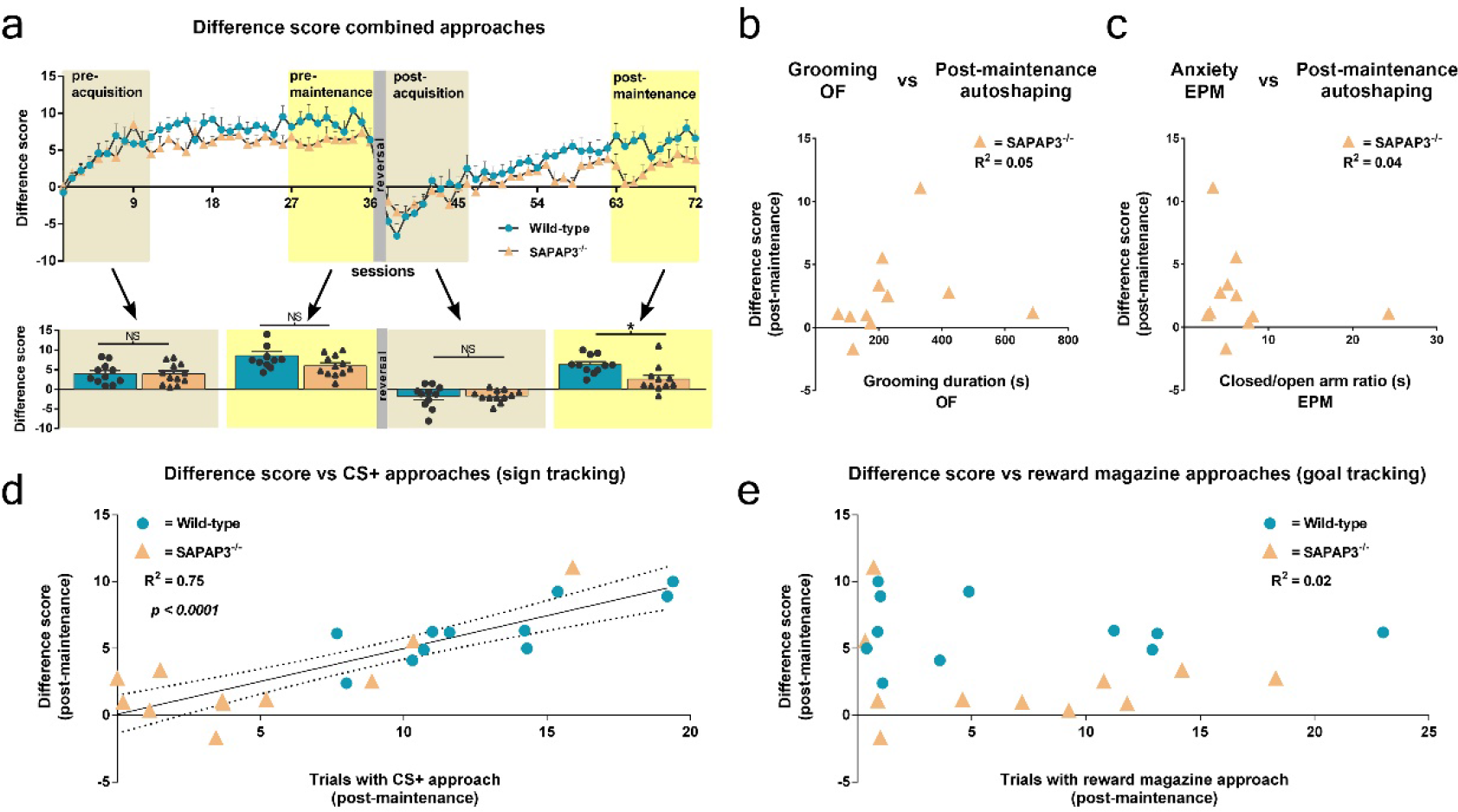
Diminished behavioral flexibility in SAPAP3^−/−^ is related to reduced sign tracking, but not compulsive grooming or anxiety. Difference score [combined CS+ approaches minus combined CS− approaches] for wild-type littermates (n=11, blue) and SAPAP3^−/−^ (n=12, orange). Bar graphs show comparison of genotypes during pre-acquisition (first 10 sessions), pre-maintenance (last 10 sessions before reversal), post-acquisition (first 10 sessions after reversal), and post-maintenance (last 10 sessions after reversal) for wild-type littermates (circles over blue bars) and SAPAP3^−/−^ (triangles over orange bars) (a). No correlation between average grooming in the open field (OF) and post-maintenance autoshaping performance (difference score) (b). No correlation between anxiety on the elevated plus maze (ratio of time spent closed/open arms) and post-maintenance performance (c). Strong correlation between CS approaches and post-maintenance performance (d). No correlation between reward magazine approaches and post-maintenance performance (e). Data are mean+SEM; * p<0.05. OF=open field; EPM=elevated plus maze; CS=conditioned stimulus.

A significant main effect of reversal on maintenance was found (F_(1,20)_=11.38, *p*=*.003*), accompanied by a significant main effect of genotype (F_(1,20)_=10.47, *p*=*.004*), without an interaction effect (F_(1,20)_=0.51, *NS*). Interestingly, post-hoc analyses revealed no significant difference between WT and SAPAP3^−/−^ before reversal (Fig 5a bar graphs pre-maintenance: 8.58±1.06 difference score WT vs. 5.98±0.75 difference score SAPAP3^−/−^, t_(21)_=2.02, *NS*); however, a significant difference after reversal (Fig. 5a bar graphs post-maintenance: 6.31±0.69 difference score WT vs. 2.58±1.02 difference score SAPAP3^−/−^, t_(20)_=3.03, *p*=*.028*) was found.

To examine the relation between grooming and autoshaping performance in SAPAP3^−/−^, correlation analyses between the grooming trait measure and the difference score were performed. No correlation between grooming and pre-maintenance difference score was found (Pearson *R^2^* of 0.004; *NS*; 95% CI −0.53-0.62), nor between grooming and post-maintenance difference score (Pearson *R^2^* of 0.05; *NS*; 95% CI −0.44-0.73) (Fig. 5b).

We explored the relation between anxiety and autoshaping performance in SAPAP3^−/−^. No correlation was found between anxiety and pre-maintenance difference score (Pearson *R^2^* of 0.08; *NS*; 95% CI −0.34-0.74), or between anxiety and post-maintenance difference score Pearson *R^2^* of 0.04; *NS*; 95% CI −0.72-0.45) (Fig. 5c).

Finally, successful reversal in WT was accompanied by re-emerging of cue approach behavior (Fig. 4c). Thus, we tested the correlation between cue approach behavior during CS+ and difference score after reversal and found a strong relationship between these two measures (Pearson *R^2^* of 0.75; *p*<*0.0001*; 95% CI 0.69-0.94) (Fig. 5d), suggesting that CS+ approach behavior is involved in successful reversal learning. No correlation was found between reward magazine approach behavior during CS+ and difference score after reversal (Pearson *R^2^* of 0.02; *NS*; 95% CI −0.53-0.30) (Fig. 5e).

## DISCUSSION

The first aim of this study was to investigate the ability of SAPAP3^−/−^, a transgenic mouse model for compulsive behavior in psychiatric disorders such as OCD, to acquire Pavlovian conditioned responding and their ability to flexibly adjust acquired behavior to reversed reward contingencies. We found that SAPAP3^−/−^, although less vigorous in their responses compared to WT, acquired responses to Pavlovian CS, but unlike WT, were unable to adapt their conditioned approach behavior upon contingency reversal. Both genotypes developed Pavlovian “goal-tracking” approaches during the CS+ (reward-magazine approaches), but ceased to goal-track after reversal. In contrast to SAPAP3^−/−^, WT exhibited “sign tracking” (CS+ approaches), which emerged during task acquisition and re-emerged towards the new CS+ after reversal, suggesting that this behavioral strategy contributed to successful reversal learning. Our second aim was to assess how behavioral flexibility is related to other OCD-like symptoms such as compulsive behavior and anxiety. Surprisingly, both grooming and anxiety-like behavior were unrelated to Pavlovian behavioral flexibility in SAPAP3^−/−^, suggesting that these traits have independent etiologies. Together, our results refine the SAPAP3^−/−^ mouse model of OCD by identifying another OCD-like trait and its relationship to other cardinal OCD-like symptoms.

SAPAP3^−/−^ have been shown to groom excessively to the point of removing fur and occasionally producing skin lesions (16). Because of these negative consequences, this behavior is considered compulsive (16). Consistently, we confirm that SAPAP3^−/−^ display increased grooming compared to WT, reflected in both number of grooming bouts and duration of grooming. Increased grooming was detected both before and after the Pavlovian conditioning and on the EPM, suggestive of a stable phenotype that is not affected by behavioral testing. Furthermore, grooming before and after autoshaping was correlated significantly, indicating that individual mice display a relatively reliable degree of grooming, even over a period of months. Our results are consistent with previous reports, demonstrating robustness of the SAPAP3^−/−^ grooming phenotype and further validate this behavioral readout as a proxy for compulsivity (16, 30).

In addition to grooming, we measured other behavioral traits that are central to OCD symptomology. We assessed anxiety on the EPM and confirmed previously reported augmentation of anxiety in SAPAP3^−/−^ (16). Previous studies measured anxiety in the OF, in the light-dark box, and on the elevated zero-maze (16). The light-dark box test and the elevated mazes are widely used assays for anxiety-like behavior (36, 37), thought to assess different forms of anxiety, bright-space and open-space anxiety, respectively (38). Thus, SAPAP3^−/−^ show increased anxiety-like behavior on different paradigms, indicating a broad anxiety phenotype.

We then asked whether increased grooming and anxiety in SAPAP3^−/−^ were related, but found no correlation between these two variables. More specifically, the degree of anxiety measured on the EPM did neither correlate with grooming on the EPM itself (grooming state during paradigm), nor with grooming repeatedly assessed in the OF (grooming trait over time). Therefore, our findings imply that compulsive behavior and anxiety are not causally related to one another in SAPAP3^−/−^, a question of clinical relevance, where some hypothesized that anxiety causes compulsion in OCD, and others hypothesized that compulsivity causes anxiety (39).

Unexpectedly, we discovered another SAPAP3^−/−^ trait that is not commonly reported as an OCD symptom: General activity (i.e., locomotion plus overall movement) was diminished compared to WT. This decreased activity was not an indirect consequence of SAPAP3^−/−^ spending more time grooming instead of being active otherwise, because decreased activity remained, even after grooming periods were excluded from the analysis (i.e., activity relative to time spent *not* grooming). In addition, this relative inactivity was not correlated with grooming itself. To ensure that this differential activity did not confound Pavlovian learning, we took several measures: 1) Animals were required to initiate trials in the autoshaping task, which enabled the exclusion of low-activity sessions and caused the total number of initiated trials not to differ between genotypes. 2) Rather than analyzing the total number of approaches during CS presentation, we analyzed the number of trials in which mice completed at least one response during the CS. 3) In order to not bias towards exclusive approaches to either cue or magazine, a measure of ‘combined approach’ responding during CS presentation was computed (i.e., counting whether an animal approached either CS or reward magazine during CS presentation). Together, these methods precluded general activity differences between genotypes from penetrating learning variables (instead of assessing how vigorous a mouse responded) and enabled direct comparison of SAPAP3^−/−^ and WT.

Although we focused on minimizing the potentially confounding effects of decreased SAPAP3^−/−^ general activity on reversal learning, it cannot be excluded as a trait of potential OCD relevance. For instance, OCD shows high comorbidity with depression and anhedonia (40–42), pathologies that produce decreased activity marked by loss of motivation and inability to experience pleasure. Furthermore, patients with severe OCD tend to exhibit depressive symptoms, elaborate avoidance behavior, and high levels of anhedonia, all of which consistent with decreased general activity. Finally, it has been reported that OCD patients move around less in their homes during everyday life compared to healthy controls (42). However, whether diminished general activity is an underexplored symptom of OCD that could potentially be studied in SAPAP3^−/−^ will have to be evaluated in future studies.

We demonstrate that SAPAP3^−/−^ were able to learn to discriminate between environmental stimuli predicting reward (CS+) and no reward (CS−) similar to WT and displayed Pavlovian conditioned approach responses during presentation of these stimuli, indicating no overall Pavlovian learning deficit. However, already during initial acquisition (prior to reversal), SAPAP3^−/−^ employed a different approach strategy than WT. In anticipation of reward, WT approached both the CS location and the reward magazine equally during CS+ presentation, whereas SAPAP3^−/−^ only approached the magazine. These two approach strategies are thought to differ in the amount of incentive salience assigned to the CS (27, 43). Approach towards the CS+ itself (sign tracking) is thought to be rooted in the CS gaining incentive salience (44), a process consistent with model-free learning (45). In contrast, approach towards the reward location (goal tracking) suggests underlying model-based learning independent of incentive motivation (46). Surprisingly, after reversal, both genotypes refrained from reward magazine approaches. SAPAP3^−/−^ did not recover responding, whereas WT re-acquired approach behavior under the reversed reward contingencies, although exclusively towards the CS+, suggesting model-free mechanisms to enable this flexible behavior. To take this speculation one step further: The lack of model-free learning-based approaches in SAPAP3^−/−^ may explain their inability to adapt to the reversal. However, future studies are necessary to test these ideas in more depth.

Previous studies indicate crucial involvement of the prefrontal cortex (PFC) in reversal learning. One PFC region, the orbitofrontal cortex (OFC), is thought to be particularly important, as OFC lesions consistently result in impaired reversal learning (47–54). SAPAP3^−/−^ display altered OFC-striatal activity (16, 17) and deficits in behavioral response inhibition that can be rescued by optogenetic stimulation of the OFC-striatal network (19). We report that once SAPAP3^−/−^ learned CS contingencies, they were unable to update their behavioral response upon reversal. One explanation for this finding is that SAPAP3^−/−^ were not able to ‘disinhibit’ responding for previously unrewarded cues, despite successful inhibition of responding to the previously rewarded cue. This is consistent with the reported intact acquisition of Pavlovian responses, but impaired reversal learning in OFC-lesioned animals (52–54). Thus, a compromised PFC-striatal network present in SAPAP3^−/−^, which was shown to be involved in their excessive grooming (16–19), is possibly responsible for the lack of adaptation to changing situational requirements. Furthermore, striatal regions that receive PFC input are thought to be critical for model-free learning (55), suggesting that both the lack of model-free response strategies in SAPAP3^−/−^ and their behavioral inflexibility may be a consequence of SAPAP3^−/−^-inherent PFC-striatal dysfunction.

The persistent, compulsive behavior of OCD patients can be conceptualized as inflexible behavior. However, previous studies examining symptom-unrelated cognitive flexibility in OCD patients yielded mixed outcomes, with some studies observing behavioral deficits in reversal learning (8, 9, 56–60), whereas others did not (61–64). However, as discussed above, deficits in reversal learning are associated with altered recruitment of fronto-striatal circuitry (suggestive of altered cognitive processing), which has been observed more consistently in OCD patients during cognitively-flexibility demanding tasks (6, 7). Moreover, a recent neuroimaging study employing Pavlovian fear conditioning found that OCD patients failed to flexibly update fear responses after reversal, despite normal acquisition of fear conditioning (65). Similarly, we found that SAPAP3^−/−^ acquire Pavlovian conditioning but fail to flexibly update their responses after contingency reversal. Thus, dysfunctional cortico-striatal circuitry in both OCD patients and SAPAP3^−/−^ may be responsible for behavioral deficits in flexibly updating conditioned responses, further validating the SAPAP3^−/−^ model of OCD.

SAPAP3^−/−^ acquired Pavlovian conditioned responding, similar to WT, but failed to flexibly update their behavior upon reversal of reward contingencies. This lack of behavioral adaptation was robust and persisted for an extended period of training after contingency reversal. Such inflexibility could potentially contribute to the persistence of compulsive behavior despite negative consequences. However, individual traits of SAPAP3^−/−^ measured here (anxiety, compulsivity, flexibility, vigor) were not linearly related to one another, suggesting at least partial independence, which may prove to be of relevance for the treatment of OCD patients. In summary, we report that in addition to compulsive behavior and augmented anxiety, SAPAP3^−/−^ display decreased vigor and cognitive deficits, thereby mapping well onto OCD symptomology. Thus, our work provides further support for the use of the SAPAP3^−/−^ model to study OCD-like behavior and its underlying neurobiology.

## Acknowledgements

We thank Dr. Matthijs Feenstra (Netherlands Institute for Neuroscience) and Dr. Nienke Vulink (Amsterdam UMC) for their insightful comments on the manuscript, Dr. Nicole Yee (Netherlands Institute for Neuroscience) for her technical assistance and input on the manuscript, Ralph Hamelink (Netherlands Institute for Neuroscience) for his practical contributions to this study, and Dr. Guoping Feng (Massachusetts Institute of Technology) for providing us with SAPAP3^−/−^.

## References

1 Abramowitz JS, Taylor S, McKay D (2009): Obsessive-compulsive disorder. Lancet. 374: 491–499.

2 Denys D (2006): Pharmacotherapy of Obsessive-compulsive Disorder and Obsessive-Compulsive Spectrum Disorders. Psychiatr Clin North Am. 29: 553–584.

3 Bokor G, Anderson PD (2014): Obsessive-compulsive disorder. J Pharm Pract. 27: 116–130.

4 Gruner P, Pittenger C (2017): Cognitive inflexibility in Obsessive-Compulsive Disorder. Neuroscience. 345: 243–255.

5 Benzina N, Mallet L, Burguière E, N’Diaye K, Pelissolo A (2016): Cognitive Dysfunction in Obsessive-Compulsive Disorder. Curr Psychiatry Rep. 18. doi:10.1007/s11920-016-0720-3.

6 Remijnse PL, Nielen MMA, van Balkom AJLM, Cath DC, van Oppen P, Uylings HBM, Veltman DJ (2006): Reduced Orbitofrontal-Striatal Activity on a Reversal Learning Task in Obsessive-Compulsive Disorder. Arch Gen Psychiatry. 63: 1225.

7 Chamberlain SR, Menzies L, Hampshire A, Suckling J, Fineberg NA, Del Campo N, et al. (2008): Orbitofrontal dysfunction in patients with obsessive-compulsive disorder and their unaffected relatives. Science (80-). 321: 421–422.

8 Valerius G, Lumpp A, Kuelz A-K, Freyer T, Voderholzer U (2008): Reversal Learning as a Neuropsychological Indicator for the Neuropathology of Obsessive Compulsive Disorder? A Behavioral Study. J Neuropsychiatry Clin Neurosci. 20: 210–218.

9 Tezcan D, Tumkaya S, Bora E (2017): Reversal learning in patients with obsessive-compulsive disorder (OCD) and their unaffected relatives: Is orbitofrontal dysfunction an endophenotype of OCD? Psychiatry Res. 252: 231–233.

10 Remijnse PL, Nielen MMA, Van Balkom AJLM, Hendriks GJ, Hoogendijk WJ, Uylings HBM, Veltman DJ (2009): Differential frontal-striatal and paralimbic activity during reversal learning in major depressive disorder and obsessive-compulsive disorder. Psychol Med. 39: 1503–1518.

11 Ting JT, Feng G (2011): Neurobiology of obsessive-compulsive disorder: Insights into neural circuitry dysfunction through mouse genetics. Curr Opin Neurobiol. 21: 842–848.

12 Milad MR, Rauch SL (2012): Obsessive-compulsive disorder: Beyond segregated cortico-striatal pathways. Trends Cogn Sci. 16: 43–51.

13 Pittenger C, Bloch MH, Williams K (2011): Glutamate abnormalities in obsessive compulsive disorder: Neurobiology, pathophysiology, and treatment. Pharmacol Ther. 132: 314–332.

14 Chakrabarty K, Bhattacharyya S, Christopher R, Khanna S (2005): Glutamatergic dysfunction in OCD. Neuropsychopharmacology. 30: 1735–1740.

15 Welch JM, Wang D, Feng G (2004): Differential mRNA Expression and Protein Localization of the SAP90/PSD-95-Associated Proteins (SAPAPs) in the Nervous System of the Mouse. J Comp Neurol. 472: 24–39.

16 Welch JM, Lu J, Rodriguiz RM, Trotta NC, Peca J, Ding JD, et al. (2007): Cortico-striatal synaptic defects and OCD-like behaviours in Sapap3-mutant mice. Nature. 448: 894–900.

17 Wan Y, Ade KK, Caffall Z, Ilcim Ozlu M, Eroglu C, Feng G, Calakos N (2014): Circuit-selective striatal synaptic dysfunction in the sapap3 knockout mouse model of obsessive-compulsive disorder. Biol Psychiatry. 75: 623–630.

18 Pinhal CM, van den Boom BJG, Santana-Kragelund F, Fellinger L, Bech P, Hamelink R, et al. (2018): Differential effects of deep-brain stimulation of the internal capsule and the striatum on excessive grooming in Sapap3 mutant mice. Biol Psychiatry. doi:10.1016/j.biopsych.2018.05.011.

19 Burguière E, Monteiro P, Feng G, Graybiel AM (2013): Optogenetic stimulation of lateral orbitofronto-striatal pathway suppresses compulsive behaviors. Science (80-). 340: 1243–1246.

20 Ahmari SE, Spellman T, Douglass NL, Kheirbek MA, Simpson HB, Deisseroth K, et al. (2013): Repeated cortico-striatal stimulation generates persistent OCD-like behavior. Science (80-). 340: 1234–1239.

21 E. E. Manning, A. Y. Dombrovski, M. M. Torregrossa SEA (2018): Impaired instrumental reversal learning is associated with increased medial prefrontal cortex activity in Sapap3 knockout mouse model of compulsive behavior. Submitt Manuscr.

22 Izquierdo A, Brigman JL, Radke AK, Rudebeck PH, Holmes A (2017): The neural basis of reversal learning: An updated perspective. Neuroscience. 345: 12–26.

23 Klanker M, Feenstra M, Denys D (2013): Dopaminergic control of cognitive flexibility in humans and animals. Front Neurosci. 7: 1–24.

24 Bussey TJ, Everitt BJ, Robbins TW (1997): Dissociable effects of cingulate and medial frontal cortex lesions on stimulus-reward learning using a novel pavlovian autoshaping procedure for the rat: Implications for the neurobiology of emotion. Behav Neurosci. 111: 908–919.

25 Mackintosh NJ (Nicholas J (1974): The psychology of animal learning. Academic Press. Retrieved April 5, 2018, from https://catalogue.nla.gov.au/Record/357768.

26 Flagel SB, Robinson TE (2017): Neurobiological basis of individual variation in stimulus-reward learning. Curr Opin Behav Sci. 13: 178–185.

27 Robinson TE, Flagel SB (2009): Dissociating the Predictive and Incentive Motivational Properties of Reward-Related Cues Through the Study of Individual Differences. Biol Psychiatry. 65: 869–873.

28 Flagel SB, Watson SJ, Robinson TE, Akil H (2007): Individual differences in the propensity to approach signals vs goals promote different adaptations in the dopamine system of rats. Psychopharmacology (Berl). 191: 599–607.

29 Kabra M, Robie AA, Rivera-Alba M, Branson S, Branson K (2013): JAABA: Interactive machine learning for automatic annotation of animal behavior. Nat Methods. 10: 64–67.

30 van den Boom BJG, Pavlidi P, Wolf CJH, Mooij AH, Willuhn I (2017): Automated classification of self-grooming in mice using open-source software. J Neurosci Methods. 289: 48–56.

31 Horner AE, Heath CJ, Hvoslef-Eide M, Kent BA, Kim CH, Nilsson SRO, et al. (2013): The touchscreen operant platform for testing learning and memory in rats and mice. Nat Protoc. 8: 1961–1984.

32 Everitt BJ, Parkinson JA, Lachenal G, Halkerston KM, Rudarakanchana N, Cardinal RN, et al. (2000): Effects of limbic corticostriatal lesions on autoshaping performance in rats. Soc Neurosci Abstr. 26: 979.

33 Cardinal RN, Parkinson JA, Lachenal G, Halkerston KM, Rudarakanchana N, Hall J, et al. (2002): Effects of selective excitotoxic lesions of the nucleus accumbens core, anterior cingulate cortex, and central nucleus of the amygdala on autoshaping performance in rats. Behav Neurosci. 116: 553–567.

34 Parkinson JA, Dalley JW, Cardinal RN, Bamford A, Fehnert B, Lachenal G, et al. (2002): Nucleus accumbens dopamine depletion impairs both acquisition and performance of appetitive Pavlovian approach behaviour: Implications for mesoaccumbens dopamine function. Behav Brain Res. 137: 149–163.

35 Wright SP (1992): Adjusted P-Values for Simultaneous Inference. Biometrics. 48: 1005.

36 Hascoët M, Bourin M (2009): The mouse light-dark box test. Neuromethods. 42: 197–223.

37 Walf AA, Frye CA (2007): The use of the elevated plus maze as an assay of anxiety-related behavior in rodents. Nat Protoc. 2: 322–328.

38 Takao K, Miyakawa T (2006): Light/dark Transition Test for Mice. J Vis Exp. 104.

39 Robbins TW, Gillan CM, Smith DG, de Wit S, Ersche KD (2012): Neurocognitive endophenotypes of impulsivity and compulsivity: Towards dimensional psychiatry. Trends Cogn Sci. 16: 81–91.

40 Ruscio AM, Stein DJ, Chiu WT, Kessler RC (2010): The epidemiology of obsessive-compulsive disorder in the National Comorbidity Survey Replication. Mol Psychiatry. 15: 53–63.

41 Diniz JB, Rosario-Campos MC, Shavitt RG, Curi M, Hounie AG, Brotto SA, Miguel EC (2004): Impact of age at onset and duration of illness on the expression of comorbidities in obsessive-compulsive disorder. J Clin Psychiatry. 65: 22–27.

42 Gershoni A, Hermesh H, Fineberg NA, Eilam D (2014): Spatial behavior reflects the mental disorder in OCD patients with and without comorbid schizophrenia. CNS Spectr. 19: 90–103.

43 Flagel SB, Robinson TE, Clark JJ, Clinton SM, Watson SJ, Seeman P, et al. (2010): An animal model of genetic vulnerability to behavioral disinhibition and responsiveness to reward-related cues: Implications for addiction. Neuropsychopharmacology. 35: 388–400.

44 Berridge KC (2004): Motivation concepts in behavioral neuroscience. Physiol Behav. 81: 179–209.

45 Morrison SE, Bamkole MA, Nicola SM (2015): Sign tracking, but not goal tracking, is resistant to outcome devaluation. Front Neurosci. 9. doi:10.3389/fnins.2015.00468.

46 Dayan P, Berridge KC (2014): Model-based and model-free Pavlovian reward learning: Revaluation, revision, and revelation. Cogn Affect Behav Neurosci. 14: 473–492.

47 Dias R, Robbins TW, Roberts AC (1996): Dissociation in prefrontal cortex of affective and attentional shifts. Nature. 380: 69–72.

48 McAlonan K, Brown VJ (2003): Orbital prefrontal cortex mediates reversal learning and not attentional set shifting in the rat. Behav Brain Res. 146: 97–103.

49 Boulougouris V, Dalley JW, Robbins TW (2007): Effects of orbitofrontal, infralimbic and prelimbic cortical lesions on serial spatial reversal learning in the rat. Behav Brain Res. 179: 219–228.

50 Chudasama Y, Robbins TW (2003): Dissociable contributions of the orbitofrontal and infralimbic cortex to pavlovian autoshaping and discrimination reversal learning: further evidence for the functional heterogeneity of the rodent frontal cortex. J Neurosci. 23: 8771–8780.

51 Chang SE (2014): Effects of orbitofrontal cortex lesions on autoshaped lever pressing and reversal learning. Behav Brain Res. 273: 52–56.

52 Burke KA, Takahashi YK, Correll J, Leon Brown P, Schoenbaum G (2009): Orbitofrontal inactivation impairs reversal of Pavlovian learning by interfering with “disinhibition” of responding for previously unrewarded cues. Eur J Neurosci. 30: 1941–1946.

53 Panayi MC, Killcross S (2018): Functional heterogeneity within the rodent lateral orbitofrontal cortex dissociates outcome devaluation and reversal learning deficits. Elife. 7. doi:10.7554/eLife.37357.

54 Tait DS, Brown VJ (2007): Difficulty overcoming learned non-reward during reversal learning in rats with ibotenic acid lesions of orbital prefrontal cortex. Ann N Y Acad Sci. 1121: 407–420.

55 Yin HH, Knowlton BJ, Balleine BW (2004): Lesions of dorsolateral striatum preserve outcome expectancy but disrupt habit formation in instrumental learning. Eur J Neurosci. 19: 181–189.

56 Lacerda ALT, Dalgalarrondo P, Caetano D, Haas GL, Camargo EE, Keshavan MS (2003): Neuropsychological performance and regional cerebral blood flow in obsessive-compulsive disorder. Prog Neuro-Psychopharmacology Biol Psychiatry. 27: 657–665.

57 Bohne A, Savage CR, Deckersbach T, Keuthen NJ, Jenike MA, Tuschen-Caffier B, Wilhelm S (2005): Visuospatial abilities, memory, and executive functioning in trichotillomania and obsessive-compulsive disorder. J Clin Exp Neuropsychol. 27: 385–399.

58 Bucci P, Galderisi S, Catapano F, Di Benedetto R, Piegari G, Mucci A, Maj M (2007): Neurocognitive indices of executive hypercontrol in obsessive-compulsive disorder. Acta Psychiatr Scand. 115: 380–387.

59 de Geus F, Denys DAJP, Sitskoorn MM, Westenberg HGM (2007): Attention and cognition in patients with obsessive?compulsive disorder. Psychiatry Clin Neurosci. 61: 45–53.

60 Cavedini P, Zorzi C, Piccinni M, Cavallini MC, Bellodi L (2010): Executive Dysfunctions in Obsessive-Compulsive Patients and Unaffected Relatives: Searching for a New Intermediate Phenotype. Biol Psychiatry. 67: 1178–1184.

61 Abbruzzese M, Ferri S, Scarone S (1997): The selective breakdown of frontal functions in patients with obsessive-compulsive disorder and in patients with schizophrenia: A double dissociation experimental finding. Neuropsychologia. 35: 907–912.

62 Cavedini P, Ferri S, Scarone S, Bellodi L (1998): Frontal lobe dysfunction in obsessive-compulsive disorder and major depression: A clinical-neuropsychological study. Psychiatry Res. 78: 21–28.

63 Moritz S, Birkner C, Kloss M, Jahn H, Hand I, Haasen C, Krausz M (2002): Executive functioning in obsessive-compulsive disorder, unipolar depression, and schizophrenia. Arch Clin Neuropsychol. 17: 477–483.

64 Fenger MM, Gade A, Adams KH, Hansen ES, Bolwig TG, Knudsen GM (2005): Cognitive deficits in obsessive-compulsive disorder on tests of frontal lobe functions. Nord J Psychiatry. 59: 39–44.

65 Apergis-Schoute AM, Gillan CM, Fineberg NA, Fernandez-Egea E, Sahakian BJ, Robbins TW (2017): Neural basis of impaired safety signaling in Obsessive Compulsive Disorder. Proc Natl Acad Sci. 114: 3216–3221.

